# Age-related increases in IGFBP2 increase melanoma cell invasion and lipid synthesis

**DOI:** 10.1101/2023.05.02.539059

**Authors:** Gretchen M. Alicea, Marie E. Portuallo, Payal Patel, Mitchell E. Fane, Alexis E. Carey, David Speicher, Hsin-Yao Tang, Andrew V. Kossenkov, Vito W. Rebecca, Denis G. Wirtz, Ashani T. Weeraratna

## Abstract

Aged melanoma patients (>65 years old) have more aggressive disease relative to young patients (<55 years old) for reasons that are not completely understood. Analysis of the young and aged secretome from human dermal fibroblasts identified >5-fold levels of insulin-like growth factor binding protein 2 (IGFBP2) in the aged fibroblast secretome. IGFBP2 functionally triggers upregulation of the PI3K-dependent fatty acid biosynthesis program in melanoma cells through increases in FASN. Melanoma cells co-cultured with aged dermal fibroblasts have higher levels of lipids relative to young dermal fibroblasts, which can be lowered by silencing IGFBP2 expression in fibroblasts, prior to treating with conditioned media. Conversely, ectopically treating melanoma cells with recombinant IGFBP2 in the presence of conditioned media from young fibroblasts, promoted lipid synthesis and accumulation in the melanoma cells. Neutralizing IGFBP2 *in vitro* reduces migration and invasion in melanoma cells, and in *vivo* studies demonstrate that neutralizing IGFBP2 in syngeneic aged mice, ablates tumor growth as well as metastasis. Conversely, ectopic treatment of young mice with IGFBP2 in young mice increases tumor growth and metastasis. Our data reveal that aged dermal fibroblasts increase melanoma cell aggressiveness through increased secretion of IGFBP2, stressing the importance of considering age when designing studies and treatment.

**Significance:** The aged microenvironment drives metastasis in melanoma cells. This study reports that IGFBP2 secretion by aged fibroblasts induces FASN in melanoma cells and drives metastasis. Neutralizing IGFBP2 decreases melanoma tumor growth and metastasis.

## Introduction

Melanoma is the deadliest form of skin cancer. Despite the advances in therapy over the past several years, melanoma remains a deadly disease due to its high metastatic potential^1,2^. Older melanoma patients display a more aggressive disease with a higher rate of metastasis to distant organs and increased resistance to targeted therapy^3^. Recent evidence from our laboratory shows that factors secreted by aged fibroblasts can drive changes in the way melanoma cells invade, metastasize, and respond to therapy^4-7^. These included secreted proteins, as well as molecules such as lipids (e.g., ceramides)^5^. Metabolism serves a key role in melanoma therapy escape as well as metastasis^8,9^, and we have shown that melanoma cells take up lipids through age-related expression of the fatty acid transporter, FATP2, in response to ceramides produced by aged fibroblasts which are then utilized to escape therapy^5^. A study by Zhang et al also showed that another FATP family member, FATP1, could be induced in melanoma cells by adipocytes, allowing them to take up lipids from those adipocytes, and become more metastatic^10^. Additionally, it has been shown that increased lipids in cancer cells are a sign of aggressive tumors and increases in lipogenic enzymes such acetyl-CoA carboxylase (ACC), fatty acid synthase (FASN), and ATP citrate lyase (ACLY) are found in almost all advanced tumors^11-14^.

To determine whether aged fibroblasts could also induce lipid synthesis as well as uptake in melanoma cells, we examined secreted factors in aged fibroblasts identified as regulators of lipid synthesis. Of these, the insulin growth factor binding protein, IGFBP2, was the most interesting candidate, as high expression of IGFBP2 has been shown to correlate with mortality^15^. IGFBP2 is part of a family of 6 IGF-binding proteins, and the second most abundant IGF family member in the human body ^16,17^. Interestingly, IGFBP2 has been linked with different aging diseases such as diabetes, insulin resistance, as well as progeria (early aging)^18,19^. High IGFBP2 expression protects against type II diabetes, regulates glucose metabolism and increases during aging and even more during premature aging^19,20^. IGFBP2 specifically can bind to insulin, IGF-1 and IGF-2, but binds with highest affinity to IGF-2; however, it plays multiple roles in cancer in an IGF-independent manner. High levels of IGFBP2 correlate with proliferation, epithelial to mesenchymal transition, invasion, and reduction of cell death in glioblastoma, prostate cancer, breast cancer, and melanoma^16,21-23^. Indeed, Li et al suggest that highly expressed IGFBP2 acts as a hub for multiple signaling pathways^24^.

In melanoma IGFBP2 plays additional roles in the activation of the EGFR-STAT3 pathway, activating PDL1, and contributing to angiogenesis^24^. Furthermore, high levels of IGFBP2 correlated with resistance to MAPK inhibitors, and may represent a marker for therapy resistance in melanoma^25^. In glioblastoma and prostate cancer, it was shown that IGFBP2 expression is negatively regulated by PTEN and positively correlated with AKT expression^26^. Loss of PTEN in melanoma may therefore increase levels of IGFBP2 and, we hypothesize, impact standard of care (BRAF/MEK inhibition) for melanoma cells. Moreover, we hypothesize that this increase in lipid synthesis through IGFBP2, promotes melanoma metastasis. Finally, IGFBP2 is also a component of the senescence associated secretory phenotype, associated with increased aggressiveness in cancer, and senescence and aging are often linked^27-29^. In the current study, we explore the expression of IGFBP2 in melanoma cells in an aged vs young microenvironment, and its subsequent impact. Taken together with our previously published work, these data add a new dimension, showing that aged fibroblasts can drive melanoma cells to not only take up lipids, but to also synthesize *de novo* lipids through pathways driven by IGFBP2.

## Materials and Methods

### Cell Culture

1205 Lu, WM164, WM793 cells were maintained in MCDB153 (Sigma)/L-15 (Cellgro) (4:1 ratio) supplemented with 2% FBS and 1.6 mM CaCl2 (tumor growth media). WM983b were maintained in DMEM supplemented with 10% FBS. YUMM1.7 murine cultured melanoma cells were maintained in DMEM F-12 (HEPES/glutamine) supplemented with 10% FBS, and 100 units per milliliter penicillin and streptomycin. Dermal fibroblast cell lines were obtained from Biobank at Coriell Institute for Medical Research. Fibroblasts were maintained in DMEM, supplemented with 10% FBS. Cell lines were cultured at 37 °C in 5% CO2 and the medium was replaced as required. Cell stocks were fingerprinted using an AmpFLSTR® Identifiler® PCR Amplification Kit from Life Technologies TM at The Wistar Institute Genomics Facility. Although it is desirable to compare the profile with the tissue or patient of origin, our cell lines were established over the course of 40 years, long before acquisition of normal control DNA was routinely performed. However, each short tandem repeat profile is compared with our internal database of over 200 melanoma cell lines, as well as control lines, such as HeLa and 293 T. Cell culture supernatants were tested for mycoplasma using a Lonza MycoAlert assay at the University of Pennsylvania Cell Center Services.

### Heatmap Analysis

The list of 91 significantly differentially secreted proteins between aged and young fibroblasts was obtained from Kaur et al, 2018 and analyzed using Ingenuity Pathway Analysis (IPA) for enrichment of specific functions and diseases. Enrichments with at least 10 affected proteins that pass p-value<10^−5^ threshold were considered significant. Individual non cell type or cancer specific functions and diseases were combined into common categories and reported along with the total category number of unique proteins and minimal p-value. These data will be made available upon request.

### Western Blot

Cell lines were plated and collected with RIPA buffer. Total protein lysate was quantified using a Pierce BCA assay kit (# 23225, Thermo Fisher Scientific) and 25µg of protein was prepared in sample buffer, boiled, and loaded into NuPAGE™ 4-12% Bis-Tris Protein Gels (#NP0321BOX, Thermo Fisher Scientific) and run at 160 volts. Proteins were then transferred onto a PVDF membrane using the iBlot system (Invitrogen), and blocked in 5% milk/TBST for 1 hour. Primary antibodies were diluted in 5% milk/TBST and incubated at 4°C overnight. The membranes were washed in TBST and probed with the corresponding HRP-conjugated secondary antibody at 0.2µg/ml. Proteins were visualized using, ECL prime (Amersham, Uppsala, Sweden) and detected using ImageQuant™ LAS 4000(GE Healthcare Life Sciences, Pittsburgh, PA).

### Antibodies

Antibodies were purchased from the following commercial vendors and used in the following dilutions for western blot: GAPDH Cell Signaling (1:10,000 2118S), HSP90 (1:10,000, Cell Signaling 4877S), IGFBP2 (1:1,000), p-AKT (1:1,000), T-AKT (1:2,000), FASN, (1:1,000) Cell signaling.

### Matrigel Invasion Assay

Matrigel-coated (invasion) 8 μm pore size translucent 24-well plate transwell chambers (BD Biosciences, San Jose, CA, USA) were used to evaluate the migration and invasion potential of melanoma cells cultured in different conditions. Briefly, 500 μL of growth medium (20% FBS) was added to the bottom of each well and a total of 2.5 × 10^4^ cells resuspended in 250 μL of aged CM, young CM in the presence or absence of rIGFBP2 or neutralizing IGFBP2 antibody (**AF674 from R&D systems)** were seeded on top.

After 18 h incubation at 37 °C, 5% CO_2_, non-invading cells were removed by wiping the upper side of the membrane, and invading cells fixed with methanol and stained with crystal violet (Sigma-Aldrich, St. Louis, MO, USA).

### Wound Healing Assay

Melanoma cells were plated to 90% confluency and wells were scratched, using a sealed pipette tip. Melanoma cells were then cultured with aged or young CM in the presence or absence of rIGFBP2 and/or neutralizing IGFBp2 antibody (**AF674 from R&D systems)**. Wells were imaged using a Nikon TE2000 inverted microscope.

### 3D-Spheroids

5,000 melanoma cells were plated in 1.5% agar and spheroids were allowed to form for 3 days. Spheroids were then embedded in rat-tail collagen (3 mg/ml Life Technologies). Spheroids were then cultured with unconditioned media, young CM, aged CM in the presence or absence of rIGFBP2 or neutralizing IGFBP2 or shIGFBP2 media from aged fibroblasts. Spheroid invasion was imaged with an Inverted microscope. For viability studies, cell viability was measured using the LIVE/DEAD® Viability/Cytotoxicity Kit (L3224, Invitrogen). Briefly, spheroids were washed with PBS and stained with calcein-AM and Ethidium homodimer-1. The dyes were diluted in PBS and 300μl of the solution was added on the spheroid wells for 1 hour at 37°C. The spheroids were washed in PBS and imaged using a Nikon TE2000 inverted microscope.

### Immunohistochemistry (IHC)

Mouse tumor and lung sections were paraffin embedded and sectioned. Paraffin embedded sections were rehydrated through a series of xylene and different concentrations of alcohol, which was followed with a rinse in water and washed in PBS. Slides were put with an antigen retrieval buffer (#3300, Vector Labs, Burlingame, CA) and steamed for 20 minutes. Slides were then blocked in a peroxide blocking buffer (#TA060H2O2Q, Thermo Scientific) for 15 minutes, followed by protein block (#TA-060-UB, Thermo Scientific) for 5 minutes and incubated with the primary antibody of interest which was prepared in antibody diluent (S0809, Dako). Slides were put in a humidified chamber at 4C overnight. Samples were washed with PBS and incubated in biotinylated anti-rabbit (#ab64256 Abcam), followed by streptavidin-HRP solution at room temperature for 20 minutes (#TS-060-HR Thermo Scientific). Samples were then washed with PBS and incubated with AEC (3-Amino-9Ethyl-I-Carboazole) chromogen for the appropriate amount of time after optimization (#TA060SA, Thermo Scientific). Slides were then washed with water and incubated in Mayer’s hematoxylin (MHS1, Sigma) for 1 minute, rinsed with water, and mounted in Aquamount (#143905, Thermo Scientific). Lungs were assessed for localization of mCherry-positive melanoma cells using a Nikon eclipse 80I digital.

### Immunofluorescence (IF)and Quantification

Samples were fixed with 4% paraformaldehyde for 15 minutes at room temperature. After samples were washed with PBS, they were incubated with BODIPY 493/503 or BODIPY 505/515 (1:3,000, Thermo Fisher Scientific), for 15 minutes at room temperature. Samples were then washed with PBS and stained with DAPI (Invitrogen, 1:5,000) for 5 minutes. After samples were washed with PBS, they were mounted in Prolong Gold antifade reagent. Lipid droplet (BODIPY stain) intensity was quantified with Adobe Photoshop software. Channels were separated and melanoma cells’ intensity was quantified across different culture conditions. Values were then compared between conditions using unpaired *t* test.

### *In vivo* Allograft Assays

All animal experiments were approved by the Institutional Animal Care and Use Committee (IACUC) of the Johns Hopkins University (Protocol M019H421: Microenvironmental Regulation of Metastasis and Therapy Resistance) and were performed in an Association for the Assessment and Accreditation of Laboratory Animal Care (AAALAC)-accredited facility. Mice were housed in a vivarium maintained at 20 ± 2 °C, 42% humidity, with a 12-h light–dark cycle with free access to food and water. The maximum tumor size allowed under this protocol was 2000 mm^3^. No tumors in our experiments exceeded this size. The young mice were utilized at 8 weeks and the aged mice at 52 weeks (Charles River). Aged male mice were single housed and young male mice were housed in groups of no more than five per cage. YUMM1.7 mCherry murine melanoma cells (2.5 × 10^5^) were injected subdermally into aged (52 weeks old) and young (6-8 weeks old) C57/BL6 mice (Charles River). Tumors were allowed to grow and treatment with rIGFBP2, neutralizing antibody or IgG control was administered. <u>rIGFBP2 experiments:</u> young mice were IP injected with rIGFBP2 in 100ul of PBS every 2 days until tumors reach 1500mm^^2^. <u>Neutralizing IGFBP2 experiments,</u> YUMM1.7 murine melanoma cells (2.5 × 10^5^) expressing mCherry were injected subdermally into aged mice. Mice were IP injected with neutralizing IGFBP2 antibody (AF797 from R&D systems) or an IgG control at a concentration of 1mg/kg every day until tumors reached 1500mm^^2^. Tumor sizes for each experiment were measured every 2 days using digital calipers, and tumor volumes were calculated using the following formula: volume = 0.5 x (length x width^2^). Mice were euthanized after 5 weeks or when a group reached 1500 mm^3^ then and tumor and lung tissue were preserved. Half of the tissue was embedded in paraffin and the other half was flash frozen and processed for protein analysis.

### shRNA, Lentiviral Production and Infection

IGFBP2 shRNA was obtained from the TRC shRNA library available at The Wistar Institute. Lentiviral production was performed according to the protocol suggested by the Broad Institute. Briefly, 293T cells were at 70% confluency and co-transfected with shRNA plasmid and the lentiviral packaging plasmids (pCMV-dR8.74psPAX2, pMD2.G for 2^nd^ generation, pMDLg/pRRE, pRSV/REV and PMD2.G for 3^rd^ generation). Appropriate empty vector and scrambled controls were created for Overexpressing and shRNA constructs respectively. Cells were transduced with lentivirus for 48 hours, allowed 48 hours to recover and then treated with appropriate antibiotic selection (puromycin) with previously established kill curves for each cell line.

### Quantitative RT-PCR

Melanoma cells were treated with young CM or aged CM and RNA was extracted using Trizol (Invitrogen) and RNeasy Mini kit (Qiagen) as protocol instructions. 1μg RNA was used to prepare cDNA using iScript DNA synthesis kit (#1708891, Bio-Rad, CA). cDNA was diluted 1:5 before use for further reactions. Each 20μl well reaction comprised of 10μL Power SYBR Green Master mix (4367659, Invitrogen), 1μL cDNA and 1μL primer

#### 1. IGFBP2

F-5’AGCCCAAGAAGCTGCGACCAC’3

R-5’CTGCCCGTTCAGAGACATCTTGC”3

#### 2. IGFBP3

F-5′ ATGCAGCGGGCGCGAC 3′

R-5′ CTACTTGCTCTGCATGCTGTAGCA 3′

#### 3. IGFBP1

F-5’TGCTGCAGAGGCAGGGAGCCC-3′ 5′-5’AGGGATCCTCTTCCCATTCCA-3

#### 4. IGFBP4

F-5’CGCCCCCAGCAGACTTCAC-3’

R-5’CTCCTCTTTTGCACCCCTCCCATTT-3’

#### 5. IGFBP5

F-5’AGATGCCTTCAGCAGAGTG-3’ R-5’ACATGCGCCTTGATGTCGTG-3’

#### 6. IGFBP6

F-5’GACCAGGAAAGAATGTGAAAGTGA-3’ R-5’GCTCTGCCAATTGACTTTCCTTAG-3’

#### 7. 18S

F-5’GAGGATGAGGTGGAACGRTGT-3’ R-5’TCT TCA GTC GCT CCA GGT CT-3’

#### 8. SREBP-1c

F-GCGCCTTGACAGGTGAAGTC R-GCCAGGGAAGTCACTGTCTTG

Final concentration used was 0.5μM. Standard curves were generated for all primers and each set of primers were normalized to an 18S Primer pair, acquired from Invitrogen, (AM1718).

### Organotypic 3D Skin Reconstructs

Organotypic 3D skin reconstructs were generated by plating, 6.4 × 10^4^ fibroblasts in each insert on top of the acellular layer (BD #355467 and Falcon #353092) and incubated for 45 minutes at 37°C in a 5% CO2 tissue culture incubator. DMEM containing 10% FBS was added to each well of the tissue culture trays and incubated for 4 days. Reconstructs were then incubated for 1 h at 37°C in HBSS containing 1% dialyzed FBS (wash media). Washing media was removed and replaced with reconstruct media I. Keratinocytes (4.17 × 10^5^) and melanoma cells (8.3 × 10^4^) were added to the inside of each insert. Media was changed every other day until day 18 when reconstructs were harvested, fixed in 10% formalin, paraffin-embedded, sectioned, and stained.

### Reverse Phase Protein Array (RPPA)

Proteins were isolated from tumor lysates and cell lines. RPPA was performed using a total of 53 antibodies. A logarithmic value was generated, reflecting the quantitation of the relative amount of each protein in each sample. Differences in relative protein loading were determined by the median protein expression for each sample across all measured proteins using data that had been normalized to the median value of each protein. The raw data were then divided by the relative-loading factor to determine load-corrected values. Logarithmic values for each protein were mean-centered to facilitate concurrent comparisons of different proteins.

### ELISA

Conditioned media from fibroblasts (3 lines per cohort) were collected after 48 hours in DMEM growth medium and filtered through a 0.45 µm low protein binding PVDF filter. Harvested CM was normalized to the fibroblast cell count and assessed for IGFBP2 using ELISA purchased from R&D system (DGB200). Serum samples were diluted 50-fold. Assay was performed as per the manufacturer’s instructions.

## Results

We revisited secretome data from dermal fibroblasts from normal healthy human donors, obtained from Coriell and derived from the Baltimore Longitudinal Study of Aging (BLSA) that showed 91 proteins significantly differentially secreted (FDR<1%) between aged and young fibroblasts^4^. The list of proteins was analyzed for enrichment of specific functions and diseases using Ingenuity Pathway Analysis (IPA). Significantly enriched categories (p-value<10^−5^) are shown in Figure A. We identified IGFBP2 as displaying increased expression in aged fibroblasts from donors aged 65 and over when compared to young fibroblasts aged 55 and younger (Figure 1A). We confirmed this by ELISA analysis of the supernatant (conditioned media(CM), 48 hours) of young (<55y) and aged (>65y) human dermal fibroblasts grown in serum-free media (Figure 1B). Furthermore, we checked the mRNA levels of all members of the IGFBP family in the young and aged dermal fibroblasts, and only IGFBP2 was highly upregulated in the aged fibroblasts compared to young fibroblasts (Supplemental Figure 1A). Using the human protein atlas, we found that older patients have a trend of higher expression of IGFBP2 compared to young patients (Supplemental Figure 1B), corresponding to a lower survival probability in older melanoma patients (Supplemental Figure 1C). Since data in the human protein atlas indicated that melanoma patients with high IGFBP2 expression have lower probability of survival, we wanted to determine if IGFBP2 secreted from aged fibroblasts could have an impact on melanoma cells. To determine whether IGFBP2 in fibroblasts could impact expression of IGFBP2 in melanoma cells, we first built skin reconstructs, an artificial 3D representation of human skin as previously described^30^. Skin reconstructs were built with either young or aged fibroblasts and identical human melanoma cell lines. We have previously shown that melanoma cells behave very differently in aged skin reconstructs, growing and invading more rapidly than when they are built with young fibroblasts^4^. When built into reconstructs with aged fibroblasts, melanoma cells increased the expression of IGFBP2 (Figure 1C, and Supplementary Figure 1D). To ascertain if this could be replicated *in vivo*, we then injected YUMM1.7 melanoma cells, derived from the BRAF/PTEN mouse model, intradermally into C57BL6 mice, of either 6 weeks of age, or >52 weeks of age. We stained the subsequent tumors with IGFBP2 antibody and found that the tumors from the aged mice stained more intensely for IGFBP2 than those from young mice (Figure 1C, Supplementary Figure 1E). Reverse phase protein array of the tumor lysates also confirmed increased IGFBP2 in melanoma in aged mice (Figure 1E), as well as an increase in fatty acid synthesis proteins, AKT and mTOR pathway (Figure 1F). Overall, our data were consistent with increases in IGFBP2 in fibroblasts during aging, and correspondingly in tumors in an aged microenvironment.

**Figure 1:**
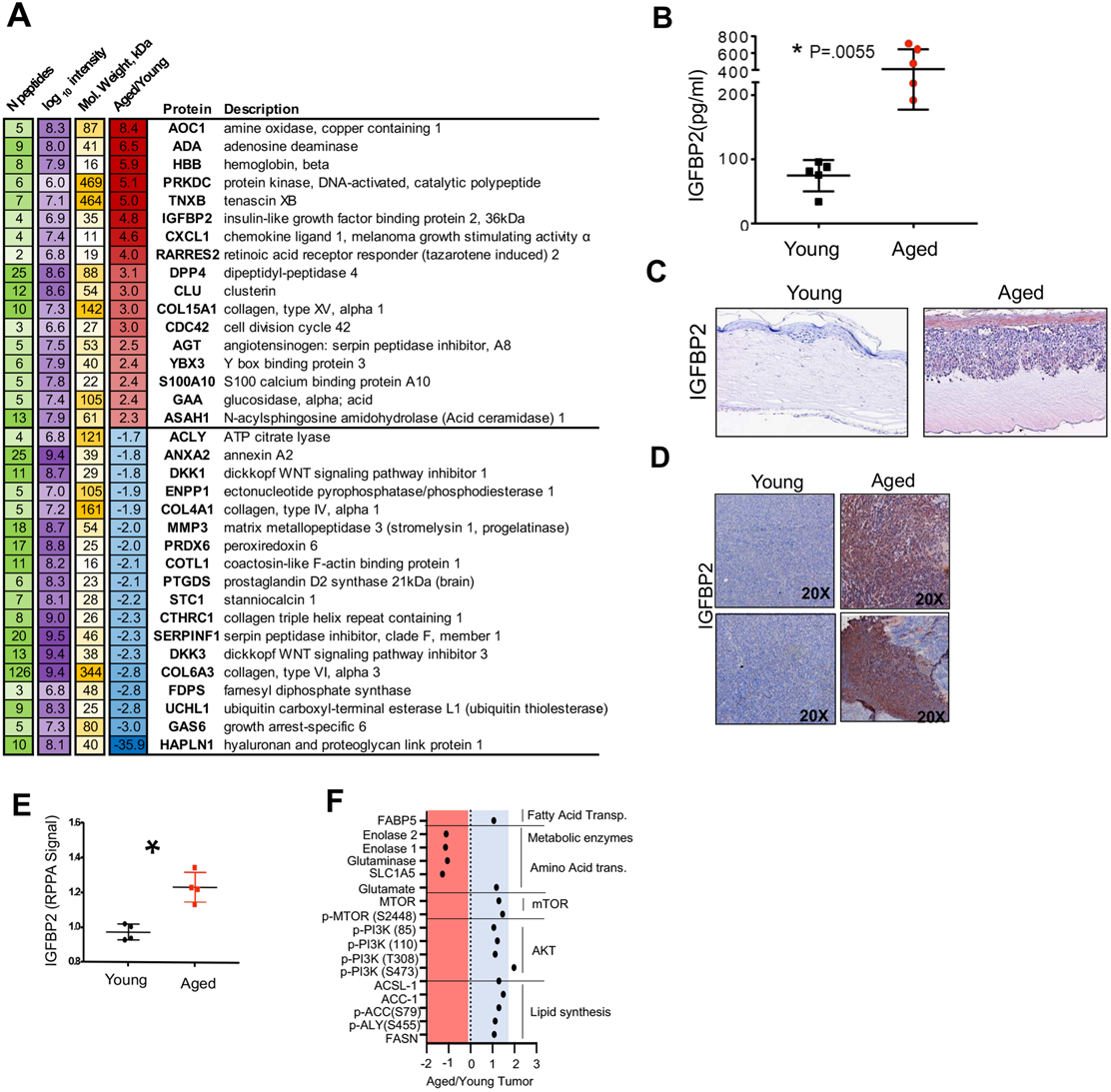
Aged fibroblasts secrete high levels of IGFBP2. Secretome analysis from conditioned media from young and aged dermal fibroblasts showing differentially overexpressed metabolic-related proteins between the two groups (Aged red/positive, young blue/negative). B) IGFBP2 ELISA analysis in young and aged dermal fibroblasts conditioned media (p=0.0055). C) IGFBP2 staining in human melanoma skin reconstructs with young or aged patient-derived dermal fibroblasts. D) IGFBP2 staining in tumor tissue from young and aged c57BL6 mice. E) RPPA analysis of young and aged Yumm1.7 mouse tumor lysate(p=0.0068). F) Pathway analysis of RPPA analysis of young and aged Yumm1.7 mouse tumor lysate.

As we observed an increase in signaling through the AKT and mTOR pathway in our RPPA data we wanted to confirm that increased IGFBP2 altered these signaling pathways in melanoma. We validated this data with the tumor lysates from our animal experiments and observed an increase in both IGFBP2 and AKT signaling (Figure 2A). Additionally, we observed a correlation of high IGFBP2 expression and high phosphorylation (S473 and T308) of AKT (Supplemental Figures 2A and 2B). To confirm that this increase in p-AKT is through IGFBP2, we cultured melanoma cells with young conditioned media and young conditioned media with recombinant IGFBP2 (rIGFBP2). We found that melanoma cells cultured in young CM and r-IGFBP2 displayed elevated levels of p-AKT (Supplemental Figure 2C). Next, we cultured melanoma cells with fibroblast conditioned media, and analyzed the fatty acid related signal transduction pathways downstream of IGFBP2 activation. We found increases in AKT and FASN in melanoma cells exposed to aged conditioned media (Figure 2B) as well as SREBP1 (Figure 2C). As we have previously published, when exposed to aged vs. young fibroblasts, melanoma cells increase lipid accumulation as demonstrated by BODIPY staining. To determine the contribution of IGFBP2 to this accumulation, we knocked down IGFBP2 in aged fibroblasts (Supplemental Figure 2D), and then exposed melanoma cells to conditioned media from the control and IGFBP2 knockdown fibroblasts. We find that knockdown of IGFBP2 in the aged fibroblasts decreases BODIPY staining in melanoma cells (Figure 2D, Supplemental Figure 2E). We also treated melanoma cells grown in young CM with recombinant IGFBP2 and showed that they increased their levels of IGFBP2, and key downstream signaling markers such as p-AKT (Figure 2E), as well as increased the levels of BODIPY staining in the melanoma cells (Figure 2F, Supplemental Figure 2F).

**Figure 2.**
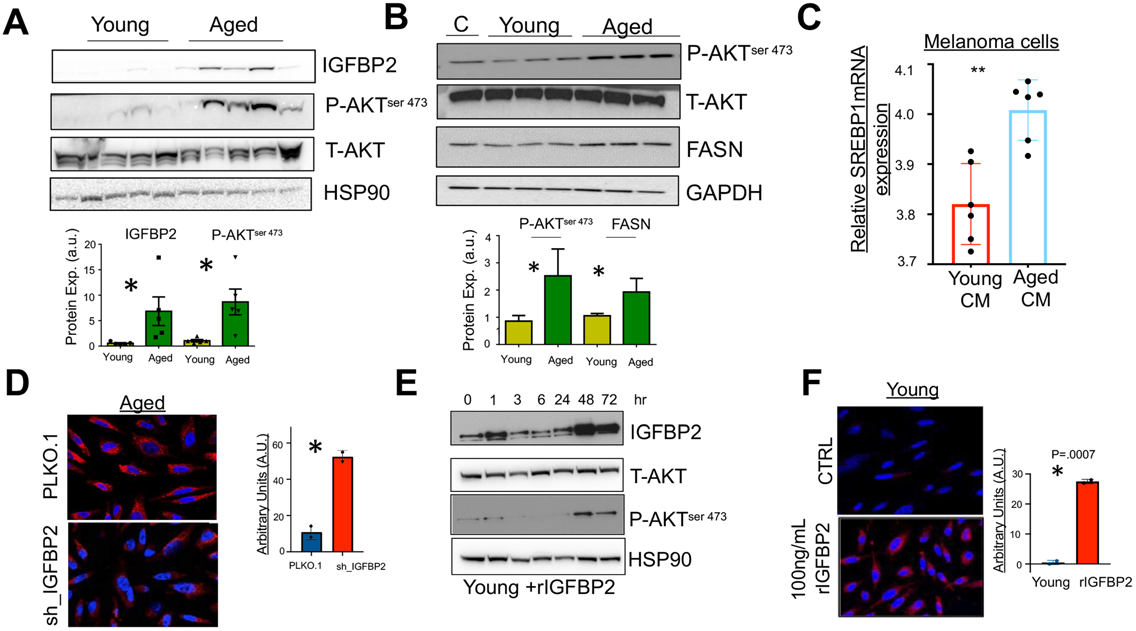
IGFBP2 induces fatty acid synthesis in melanoma cells. A) Western blot analysis of IGFBP2, phospho-AKT, total AKT, and HSP90 (loading control) from young and aged tumor lysate. Quantification of immunoblotting of IGFBP2 and phospho-AKT relative to HSP90 loading control. B) Western blot analysis of FASN, phospho-AKT, total AKT, and GAPDH (loading control) of melanoma cells (1205Lu) cultured with young and aged conditioned media. Quantification of immunoblotting of FASN and phospho-AKT relative to GAPDH loading control. C) RT-PCR analysis was performed for SREBP1 on melanoma cells (WM793) cultured with young and aged cm. (P=.0071) D) IF microscopy of human melanoma cells (1205Lu) cultures with aged fibroblasts conditioned media with PLKO.1 or shIGFBP2. Quantification of BODIPY (P=.0087) E) Western blot anslysis of melanoma cells (1205Lu) treated with recombinant IGFBP2 (150ng/ul) at different times. Cells were probed for IGFBP2, phospho-AKT, total-AKT, and HSP90 (loading control). Quantification of immunoblotting of IGFBP2 and phospho-AKT relative to HSP90 loading control. F) IF microscopy of melanoma cells treated with recombinant IGFBP2 (150ng/ul) and young conditioned media. Quantification of BODIPY (P=.0007). * Indicates p<0.05 student t-test was used.

Previous data from our laboratory has indicated that melanoma cells increase their invasion in response to CM from aged fibroblasts^4^. Since IGFBP2 increased FASN in melanoma cells in an aged microenvironment, we next wanted to observe if lipids would promote invasion in the melanoma cells. We show here that adding lipids, such as Albumax ®, a lipid-rich cocktail, can significantly increase invasion of melanoma cells (Supplemental Figure 3A). To test whether IGFBP2 in an aged vs young environment played a role in melanoma cell invasion, we first performed 2D scratch assays, followed by Boyden chamber invasion assays, 3D spheroid assays and finally *in vivo* assessments. Wound healing assay analysis showed us that aged fibroblast CM increased melanoma cell migration over young CM, and that this could be reversed using a neutralizing antibody against IGFBP2. Similarly, the impact of young CM on melanoma cell invasion could be increased using a recombinant IGFBP2 (Figure 3A, Supplemental figure 3B). We then performed a Boyden Chamber assay to test for invasion, and observed a similar result (Figure 3B), and finally performed a 3D spheroid assay, where we embedded melanoma cell spheroids in collagen with young or aged fibroblasts, in which IGFBP2 was modulated, ie., rIGFBP2 in young fibroblasts (Figure 3C, Supplemental Figure 3C) or shIGFBP2 in aged fibroblasts (Figure 3C, Supplemental Figure 3D). In all of these experiments, aged fibroblasts were able to increase invasion of melanoma cells, but less so in the absence of IGFBP2, while rIGFBP2 was able to increase invasion of melanoma cells in both young CM, or in the presence of young fibroblasts.

**Figure 3.**
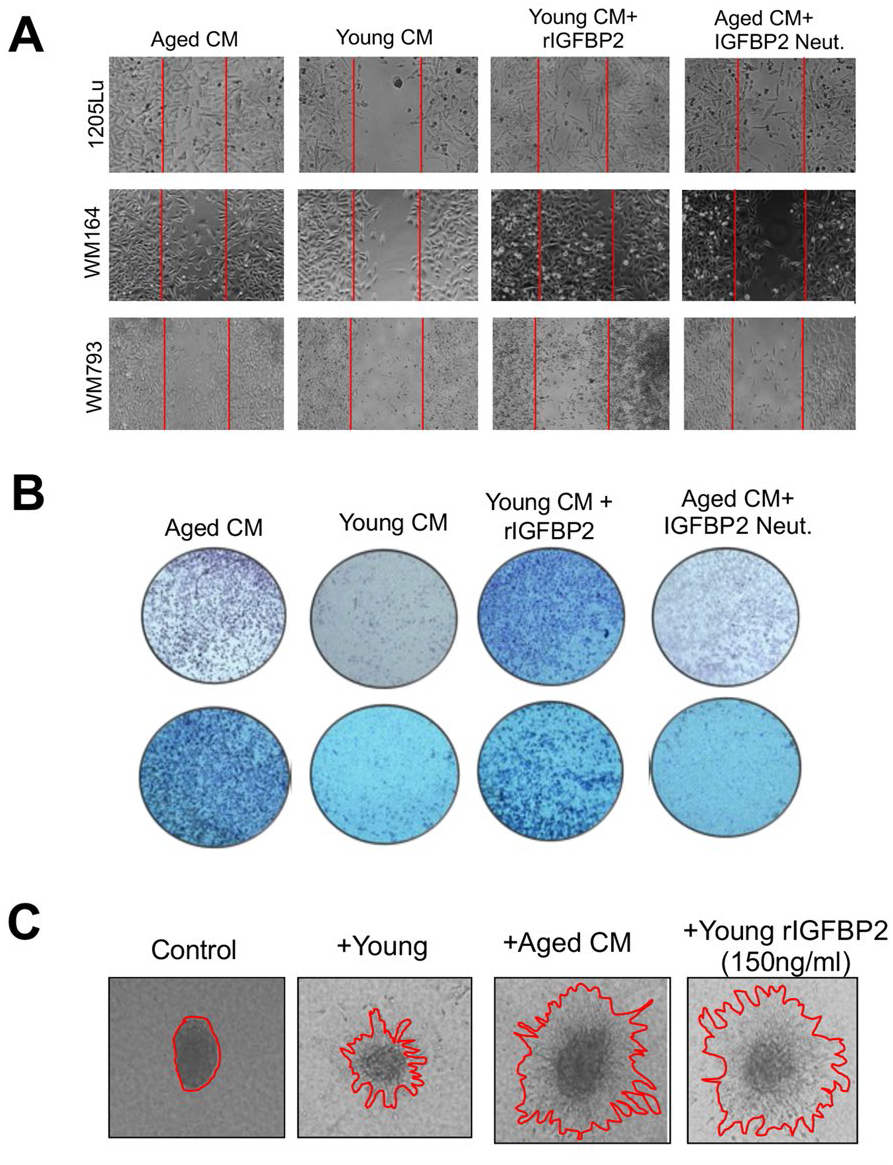
IGFBP2 increases melanoma cell migration and invasion. **A**) Wound healing assay of human melanoma cells (1205Lu, WM164, WM793) cultured with young, aged conditioned media in the presence or absence of a neutralizing IGFBP2 antibody (10uM) or recombinant IGFBP2. (150ng/ul) B) Matrigel invasion assay of melanoma cells (1205Lu, WM164) cultured with young and aged conditioned media in the presence or absence of a neutralizing IGFBP2 antibody or recombinant IGFBP2. C) Melanoma cells (1205Lu) grown in 3D spheroids cultured with aged CM, and young CM in the presence or absence of recombinant IGFBP2 (150ng/ul) for 48 hrs.

Finally, we wanted to test the impact of modulating IGFBP2 in young and aged mice in vivo. We first examined the impact of IGFBP2 modulation on the growth of mCherry tagged YUMM 1.7-melanoma cells in vivo. As observed in other cancers, recombinant IGFBP2 significantly increased the growth of tumors in young mice (Figure 4A). To confirm that the tumors have high expression of IGFBP2, we performed IHC and confirmed high IGFBP2 expression in the tumors of young mice treated with rIGFBP2 (Figure 4B, Supplemental Figure 4A). When tumor lysates were examined by Western analysis for downstream markers of IGFBP2 signaling, rIGFBP2 increased both IGFBP2 levels and PO4-AKT in the tumors of young mice (Figure 4C). Similarly, treatment with a neutralizing antibody against IGFBP2 slowed down growth in aged mice (Figure 4E). Additionally, western analysis of tumor lysates from aged mice showed decreases in IGFBP2 and PO4-AKT levels (Figure 4F). To test if IGFBP2 can also promote metastasis, we isolated the lungs of young mice and observed an increase in lung colonization upon treatment with rIGFBP2 (Figure 4D, Supplemental Figure 4B), whereas neutralizing IGFBP2 in aged mice decreased metastasis (Figure 4G, Supplemental Figure 4C) as seen by mCherry staining. Overall, these data show that IGFBP2 plays a role in driving melanoma metastasis and growth, in an age-specific manner, and neutralizing IGFBP2 levels in mice decreases tumor growth and lung colonization.

**Figure 4.**
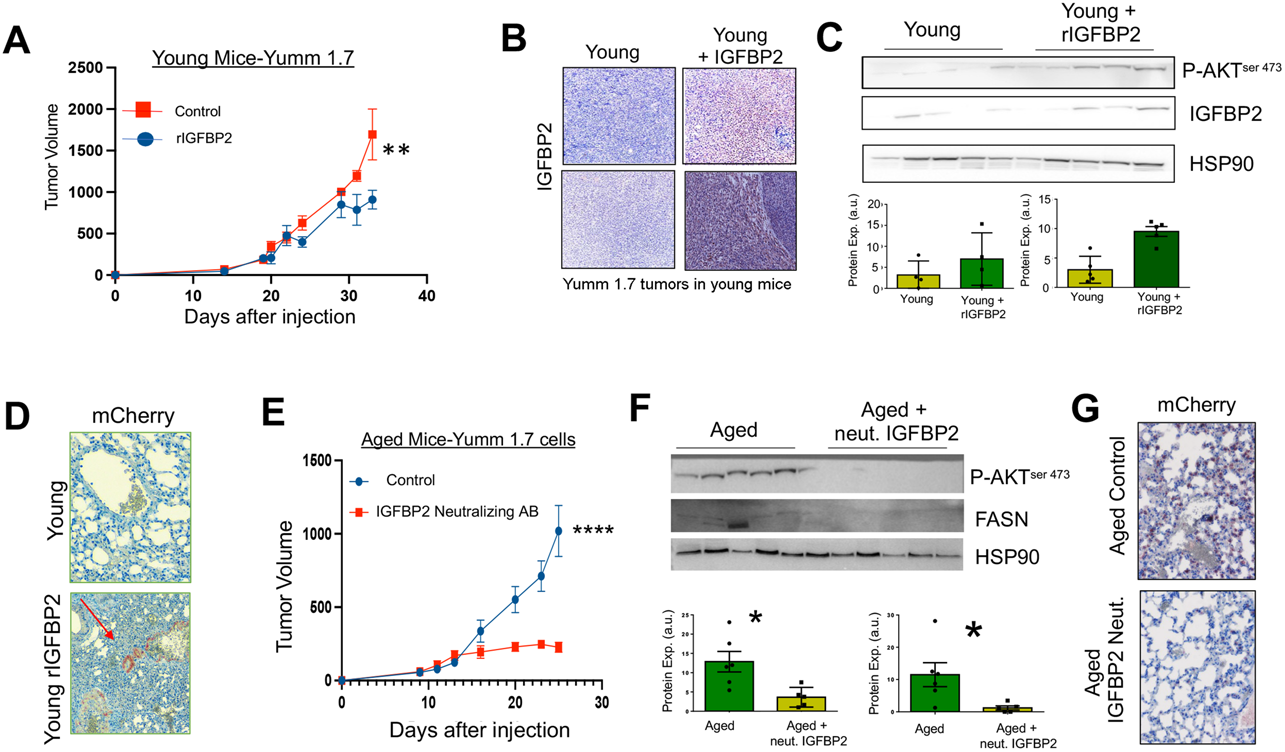
IGFBP2 increases melanoma tumor growth and metastasis *in vivo*. A) YUMM1.7-mcherry murine melanoma cells were grown in young (8 weeks old) mice. Tumor growth of young mice after treatment with 500ng recombinant IGFBP2 or PBS (3 times a week, after tumors were palpable) were measured. B) Tumors from young mice treated with PBS or recombinant IGFBP2 underwent IHC analysis for IGFBP2. C) Protein expression analysis was performed on tumor lysate from young mice treated with PBS and recombinant IGFBP2. Densitometry of phospho-AKT and IGFBP2 immunoblotting relative to HSP90 loading control. D) Lungs from young Yumm1.7 tumor-bearing mice treated with PBS or 500ng of recombinant IGFBP2 (3 times a week) underwent IHC analysis for mCherry localization. E) Tumor growth curve of YUMM1.7 melanoma cells subdermally injected in old (52 weeks old) mice treated with IGFBP2 neutralizing antibody (at a concentration of 1mg/kg every day, n=5) vs an IgG control (n=5). F) Protein expression analysis was performed on tumor lysate from aged mice treated with IgG or neutralizing IGFBP2 antibody. Quantification analysis of phospho-AKT and FASN immunoblotting relative to HSP90 loading control G) Lungs from YUMM1.7 tumor-bearing aged mice treated with IgG or neutralizing IGFBP2 underwent IHC analysis for mCherry localization. * Indicates p<0.05 student t-test was used.

## Discussion

Older melanoma patients display more aggressive disease relative to their younger counterparts, as evidenced by a higher metastatic burden to distant organs and worse survival outcomes. Despite the advances in targeted- and immune-based melanoma treatment strategies, most elderly patients still succumb to their disease due to their metastatic burden. Immense metabolic plasticity is requisite to allow cancer cells to survive the nutritionally diverse environments they encounter as they locally invade, enter the circulation, and ultimately leave the circulation to colonize distant organs^13,31,32^. Notably, the metabolic landscape that tumor cells encounter radically changes during organismal aging, which has significant implications on tumor biology and phenotypes^33,34^. Our previous data demonstrate that host stromal fibroblasts exhibit marked metabolic reprogramming and alterations in their lipid secretion profile during organismal aging, translating to adaptive changes in melanoma cell uptake and utilization of aged fibroblast-derived lipids in the aged TME to escape therapy^5^. In agreement, it was recently reported that serum isolated from elderly patients containing by-products of methylmalonic acid (MMA) conferred chemoresistance in human triple negative breast cancer and lung cancer^34^. Nonetheless, our understanding of the dynamic relationship between organismal aging, metabolism, and tumor biology is only in its infancy.

Here, we report that in addition to elevated secretion of lipids capable of altering melanoma metabolism, aged stromal fibroblasts also secreted increased IGFBP2 levels that metabolically reprogram melanoma cells to synthesize lipid droplets that drive a pro-invasive/metastatic cell state. Lipid droplets serve essential physiological roles in cell biology, signaling, membrane formation, and energy source; and evidence shows that tumor cells also leverage lipids to drive aggressive phenotypes^35,36^. Lipid droplets are composed mainly of triglycerides, cholesterol ester surrounded with a phospholipid monolayer which plays a role not only as a source of energy but in cell membrane formation and signaling, which are used by cells to proliferate, create cell protrusions and invade^37-39^. Other studies have shown that melanoma, breast cancer and ovarian cells among others, can uptake lipids from adipocytes to increase their proliferation and invasion^10,40,41^. Our data is consistent with recent reports in other cancers that reveal the importance of IGFBP2 in tumor growth and metastasis and represents the seminal report of aged stromal fibroblasts as a significant source of IGFBP2 in the aged TME.

In line with what has been observed in the senescence field, where senescent cells secrete high levels of IGFBPs, we observe an increase in the secretion of IGFBP2 in aged fibroblasts^42,43^. The role of IGFBPs in tumor progression and therapy resistance varies by cancer type. IGFBP2 over-expression occurs in advanced cancers including ovarian cancer, prostate cancer, and glioblastoma, and its high expression has been linked to an aggressive phenotype^44-46^. Our data show that aged fibroblasts secrete elevated levels of IGFBP2 which elevates the metastatic capacity of melanoma cells. We have demonstrated that IGFBP2 from the aged fibroblasts promotes melanoma invasion and metastasis of melanoma cells. Supplementing melanoma cells with rIGFBP2 in the young microenvironment promotes melanoma invasiveness and metastasis.

Mechanistically, our studies suggest that IGFBP2 could promote melanoma metastatic capacity by activating PI3K, which leads to FASN and elevated lipid synthesis by melanoma cells. Exogenous administration of rIGFBP2 *in vivo*, increased p-AKT levels in tumors in planted in young mice. Notably, treating aged mice with IGBP2 neutralizing antibody decreases p-AKT ^ser 473^ and FASN expression. Moreover, treating young mice with r IGFBP2 increased tumor growth, while neutralizing IGFBP2 not only decreased tumor growth and lung colonization as well. Our study demonstrates the importance of the aged TME in driving tumor dissemination and highlights the need for therapeutic interventions guided by the age of the patients that prevents tumor metastasis. These data provide the rationale that IGFBP2 may serve as a tractable therapeutic target to address elevated metastatic burden of aged patients with melanoma.

## Acknowledgments

We thank the outstanding Core Facilities of the Johns Hopkins University (P30CA00697356). A.T.W., M.E.F. and A.E.C are supported by P01CA114046 and R01CA207935. A.T.W. is also supported by a Team Science Award from the Melanoma Research Alliance. A.T.W. is also supported by U01CA227550, R01CA232256, a Bloomberg Distinguished Professorship, and the EV McCollum Endowed Chair of Biochemistry and Molecular Biology. G.M.A is supported by the T32 training program in nanotechnology. D.W is supported through grants from the National Cancer Institute (U54CA143868) and the National Institute on Aging (U01AG060903).

